# Bioinformatics analysis in identifying significantly expressed immune related genes in *influenza A virus* infection

**DOI:** 10.1101/2022.08.15.503998

**Authors:** Ayinde Olaniyi, Alabi Oluwabunmi, Oguntoye Oluwatobi

## Abstract

This study is done to identify significantly expressed genes of influenza A virus infection in human, facilitated by the idea of checking for the presence of immune related genes in these significantly expressed genes, followed by the analysis of their potential mechanism and pathway. Considering the usage of microarray dataset, the gene expression profile GSE66597 was downloaded from the gene expression omnibus(GEO) database, and then a subsequent generation of differentially expressed genes(DEGs) using the GEO2R tool. A total of 1107 genes was generated with 703 upregulated and 404 downregulated genes. The protein-protein interaction (PPI) network was constructed using string and Cytoscape software which resulted in generating the core genes, hub genes, and bottleneck genes. Subsequent Venn diagram analysis finally gave the six candidate genes CRP, CCL5, IL17A, STAT1, CD34, SPI1, subjected to gene ontology (GO) and Kyoto encyclopedia of genes and genomes (KEGG) for enrichment and pathway analysis respectively. Out of the initial six candidate genes, four of these genes CRP, CCL5, IL17A, are immune related, and initiated particularly by the presence of influenza A virus.

## 1. Introduction

The continual emergence and re-emergence of viruses put man on its toes in finding long lasting solution to this menace that pose existential threat man. Severe acute respiratory tract infections (SARI) is the fourth most common cause of death worldwide as captured by the WHO [1], and respiratory viruses are the main etiological agents of SARI [2]. The influenza A virus (IAV) is the most prevalent agent among these viruses as the first breakout of IAV in 1918 reportedly claimed around 50 million lives worldwide, and till today the virus is still seen as a great threat as there have been yearly seasonal outbreaks and epidemic causing up to 650 thousand deaths each season [3].

The first breakout of IVA in 1918 reportedly claimed around 50 million lives worldwide, and till today the virus is still seen as a great threat as there have been yearly seasonal outbreaks and epidemic [4]. The virulence nature and the cause of IAV can be tied to its mechanism of infection and action as it targets particularly the respiratory tract of its host, causing chronic damages to the alveoli, hence, its characteristic fatality[5].

Due to the epidemic and pandemic potentials of this virus, novel strategies for the development of interventions especially in the form therapeutic agents are needed to be employed in combating its threat [6], more especially the development and utilizing of compounds that enhances the ability of the innate immune system to combat viral infection. Hence, the need for a targeted genetic profiling of this virus, identifying significantly expressed immune related genes, useful as a potential candidate drug target, and as early diagnostic markers. However, As reported by several studies, the instrumentality of gene expression profiling has helped to identify a number of differentially expressed genes (DEGs) involved in multiple signaling pathways, molecular functions, and biological processes, which are having an important role in the incidence and development of diseases and could be used as a potential molecular target and diagnostic markers [7]. Furthermore, it is now understood that gene expression analysis based on microarray technology is a powerful and high-throughput research method, and this can be greatly useful even in the gene expression profiling of IVA miRNAs. This method is employed in this study different from the currently and mainly analyzed IVA’s single nucleotide polymorphisms (SNPs).

In this work, we used bioinformatics tools to identify the top hub genes and bottleneck genes of the IVA by downloading GSE66597 dataset [8] from the Gene Expression Omni-bus (GEO) database to identify IVA-related DEGs between IVA(H5N1) and control samples. Furtherly, Gene Ontology (GO) enrichment analysis, Kyoto Encyclopedia of Genes and Genomes(KEGG) pathway analysis, and Protein-Protein Interaction (PPI) network analysis were performed to discover potential genes as IVA diagnostic bio-markers and therapeutic targets, as well as for further studies.

## 2. Materials and methods

### 2.1. Retrieval of microarray data

The microarray gene expression dataset GSE66597 based on the GPL6480 platform (Agilent-014850 Whole Human Genome Microarray 4×44K G4112F) was retrieved from the gene expression omnibus (GEO) database of NCBI (https://www.ncbi.nlm.nih.gov/geo/) by using the keyword multiple influenza virus. The gene expression profile contains a total of 18 samples containing 9 test samples and 9 control samples were selected and analyzed.

### 2.2. Identification of DEGs

Significant DEGs between test and control samples were analyzed by using an online analysis tool GEO2R (https://www.ncbi.nlm.nih.gov/geo/geo2r/) for the dataset. The GEO2R tool is an interactive web tool for comparing two sets of data under the same experimental conditions and can analyze any geo series [9], it uses Bioconductor packages such as GEOQuery and limma for processing of data [10]. DEGs between IVA and normal blood samples were screened with the following threshold criteria; P-value < 0.05 and a fold change value∣logFC∣≥ 1.0.

### 2.3. Integration of the protein-protein interaction (PPI) network

The PPI network of DEGs was constructed using the online-based tool STRING (https://string-db.org/) for protein-protein interaction(PPI),with confidence score of ≥ 0.9,hiding all disconnected nodes in the network.

### 2.4. Identification of core genes, hub genes and bottle-neck genes

The constructed PPI networks were visualized using-based Cytoscape software (3.7.1) (https://cytoscape.org/). MCODE plug-in of cytoscape was used to screen out the core genes that constitute the stable structure of the PPI network with default degree cut-off =3, haircut on, node score cut-off=0.2, k-core=4, and maximum depth =100. The CentiScape plug-in was used to calculate the centrality index and topological properties for the identification of the most important nodes of a network, including undirected, directed, and weighted networks. The top 15 genes termed (hub genes) and the bottleneck genes were identified using the CytoHubba plugin of cytoscape defined with degree value method, and betweenness value method respectively.

### 2.5. Functional and pathway enrichment analysis of DEGs

For the enrichment analysis, the gene ontology (GO); mainly including biological process, cellular component and molecular function, and the Kyoto Encyclopedia of Genes and Genomes (KEGG) analyses were performed by uploading candidate genes into the online-based tool Enrichr (https://maayanlab.cloud/Enrichr/). GO is a common way to annotate genes, their products, and their sequences, while the KEGG is a detailed data-base resource for the biological interpretation of genomic sequences and other high-throughput data.

### 2.6. microRNA network analysis

The candidate genes were submitted to the online software miRNet (https://www.mirnet.ca/) to find out the interaction between miRNAs and genes. A network between genes and miRNAs were generated by using the software.

## 3 Results

### 3.1. Identification of DEGs

Based on the set cut-off criteria (∣logFC∣≥2 and P < 0 .05), The differential expression analysis came out with a total of 1107 DEGs. Including 703 upregulated DEGs and 404 downregulated DEGs using the GEO2R.

### 3.2. Analysis of modules of core genes, hub genes and bottleneck genes

After hiding the un-interacting nodes, the PPI network showed a number of 570 nodes and 1295 edges involved in the network. (Figure 2(a)). MCODE K kernel analysis of the string network generated a total of 14 clusters totaling 90 core genes screened (Figures 2(b)–2(o)).; with default parameters degree cut-off = 3, haircut on, node score cut-off = 0.2, k-core = 4, and maximum depth = 15 bottleneck genes were and 15 hub genes were calculated and obtained using Cytohubba (Figure 3(a)(b)). Furthermore, using the desktop-based software FUNRICH Venn diagram analysis of the above three datasets, six candidate genes were obtained for further analysis, this includes signal transducer and activator of transcription 1 (STAT1), C-C motif chemokine ligand 5 (CCL5), interleukin 17A (IL17A), Spi-1 proto-oncogene (SPI1), CD34 molecule (CD34), and C-reactive protein (CRP) (Figure 4).

**Figure 1.**
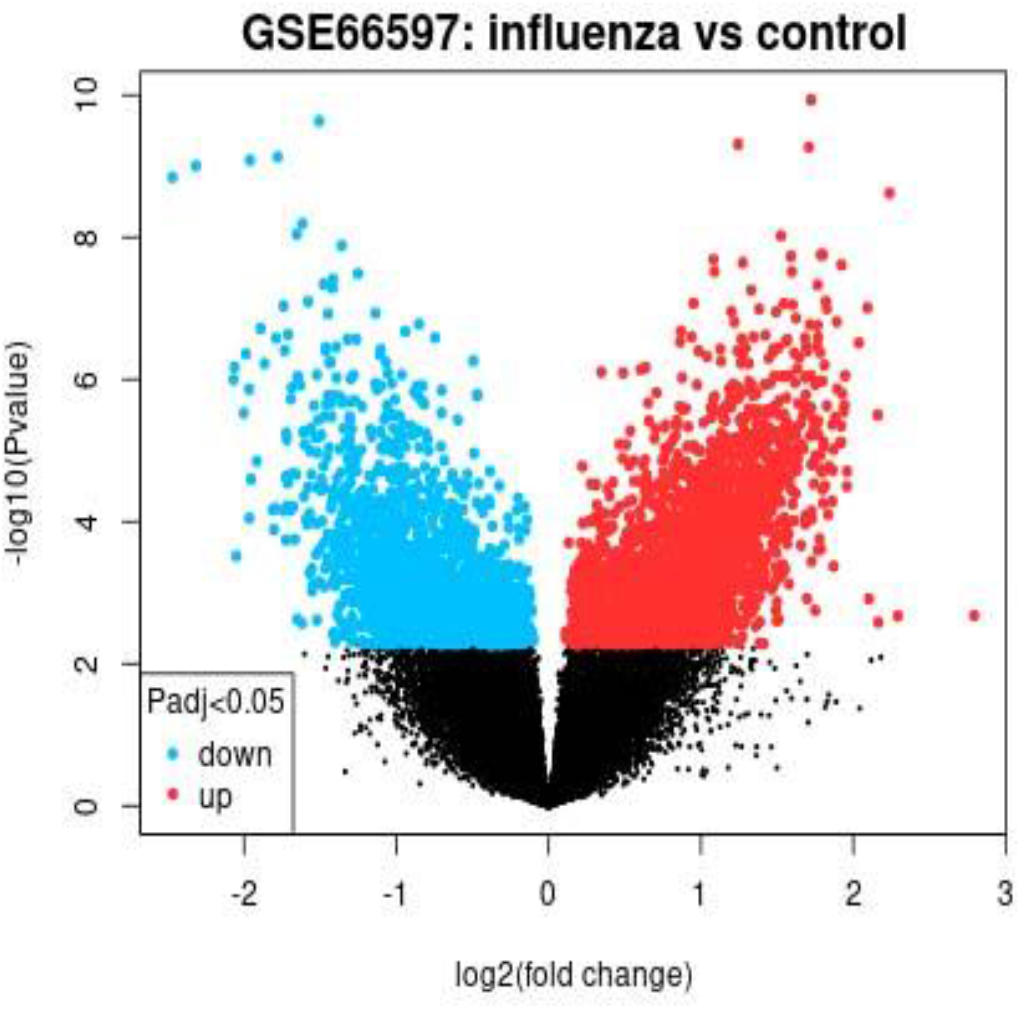
Volcano plot analysis of DEGs. Red indicates that expression of the gene is relatively upregulated while blue indicates that expression of the gene is relatively downregulated; black indicates no significant changes in gene expression.

**Figure 2.**
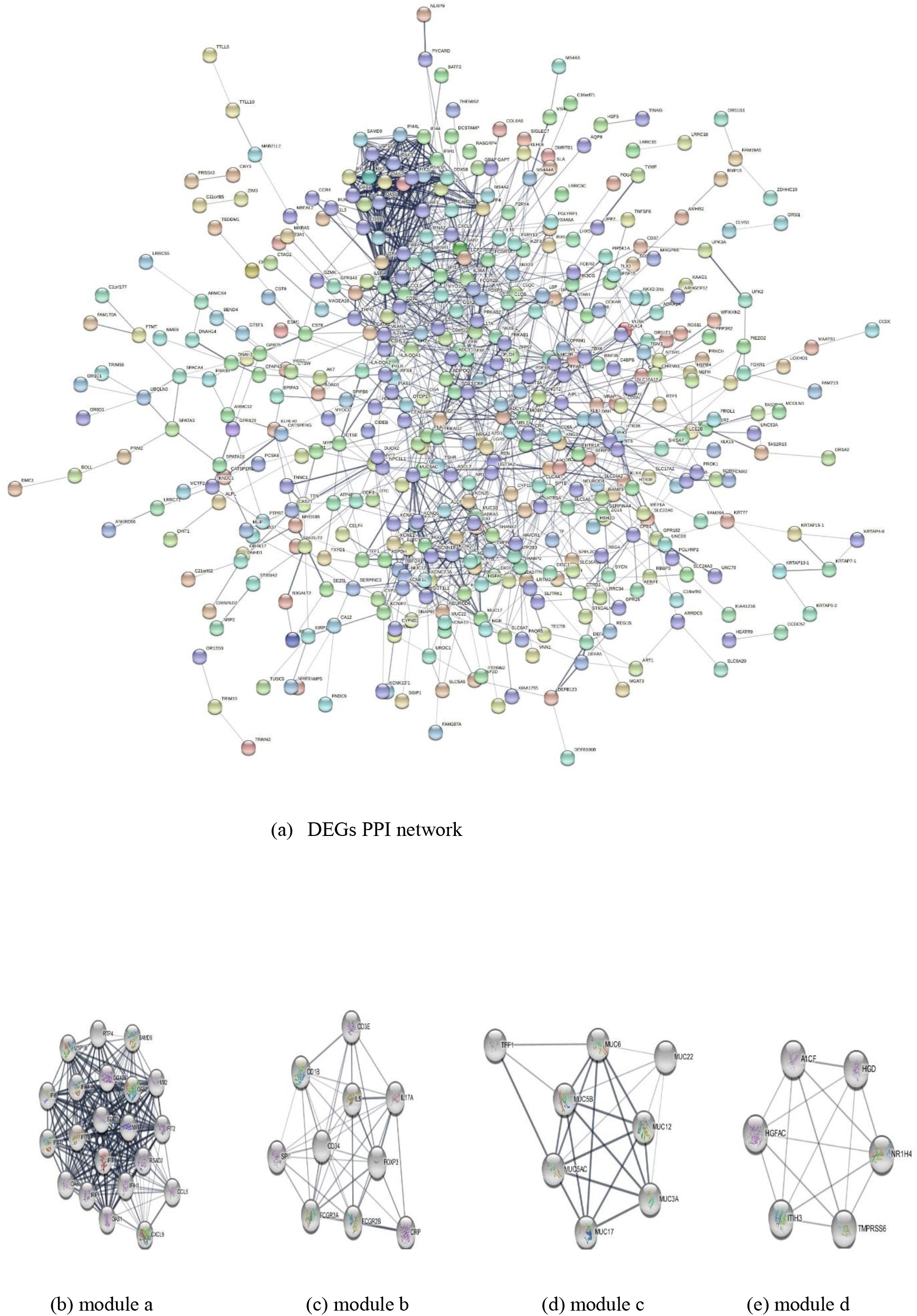
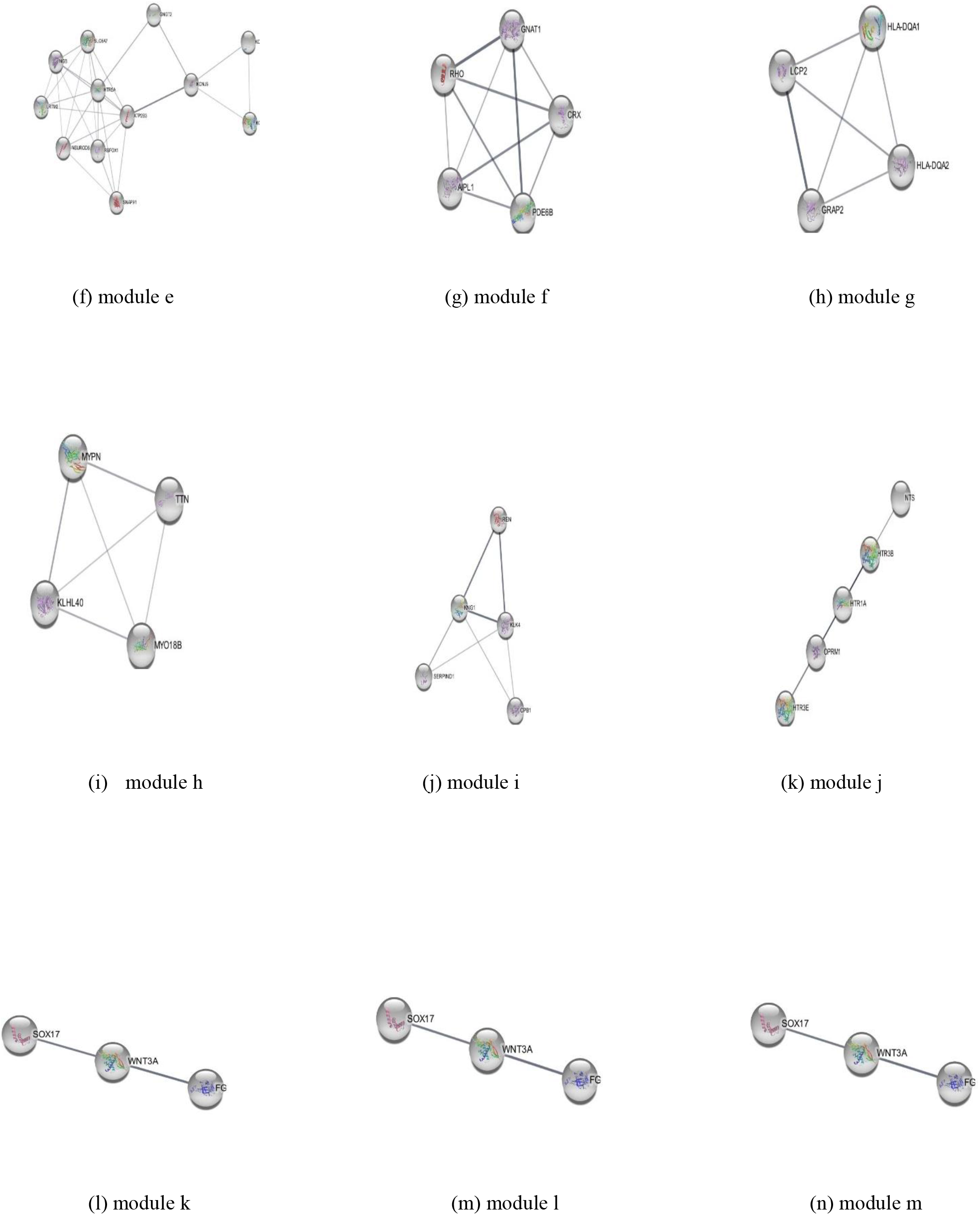
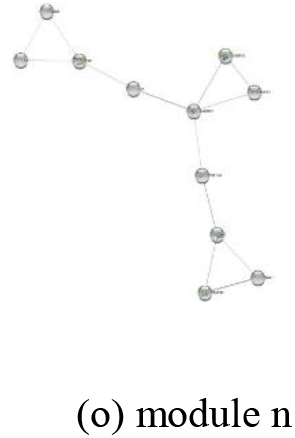
Protein-protein interaction (PPI) networks constructed by STRING tool and modular analysis

**Figure 3.**
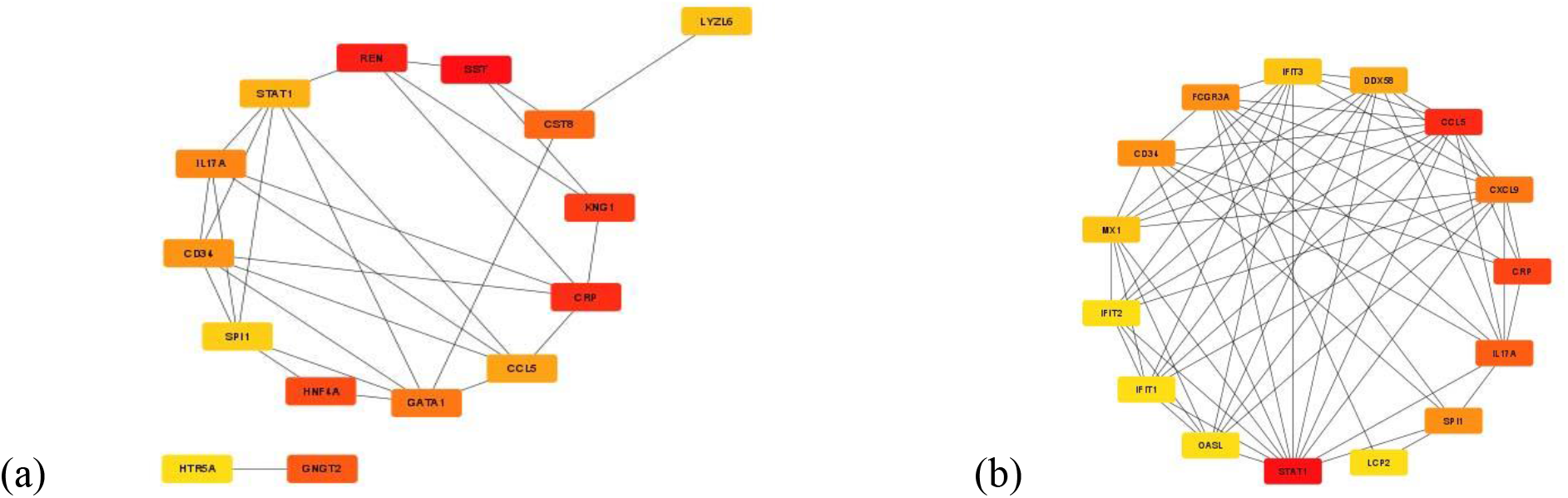
(a) Network of 15 bottleneck genes defined in degree value. (b) network of 15 hub genes defined with betweenness value. The represents the degree/betweenness of connectivity. the red color represents the highest degree/betweenness, the orange color represents intermediate degree/betweenness, and the yellow represents the lowest degree/betweenness

**Figure 4.**
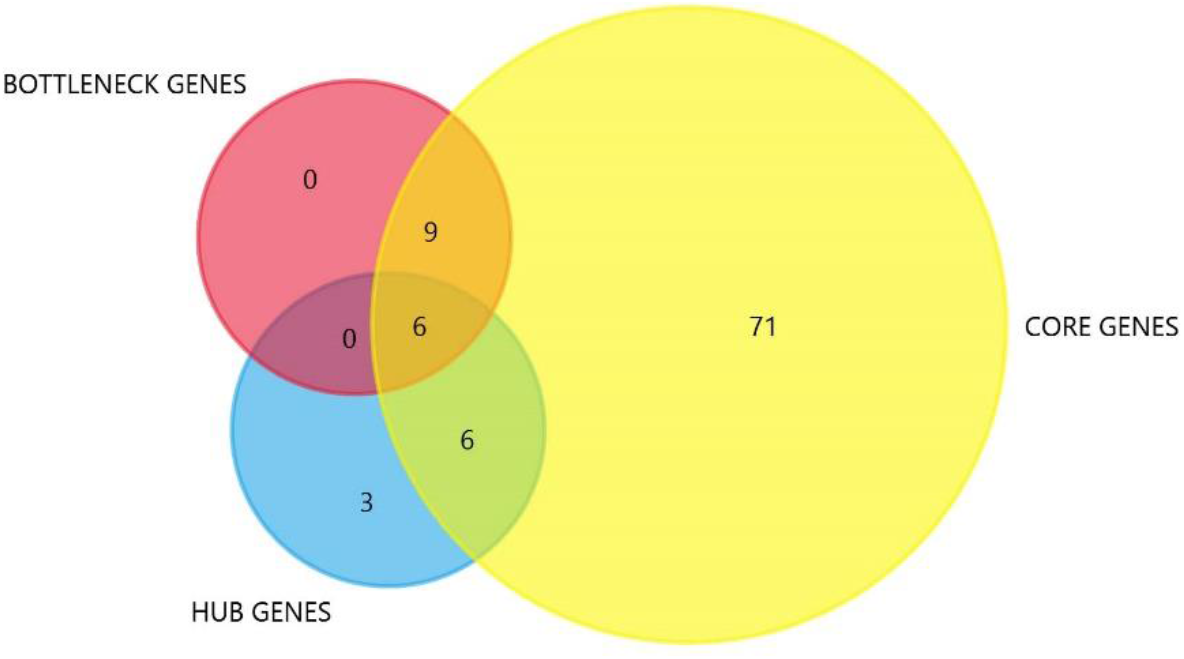
Venn diagram analysis of candidate genes. The yellow circle represents core genes, the blue circle represents hub genes, and the pink circle represents bottleneck genes

### 3.3. Functional and pathway enrichment analysis of DEGs

Three gene ontology(GO) results were obtained and presented in (Figure 5(a)-(c)). Candidate genes were significantly enriched in interleukin-6-mediated signaling pathway, cytokine-mediated signaling pathway, and cellular response to interleukin-6 considering the biological processes. For the molecular function, RNA polymerase II-specific DNA-binding transcription factor binding, CCR1 chemokine receptor binding were significantly enriched. Also, for the cellular component, candidate genes are significantly enriched in axon, lytic vacuole and dendrite. The KEGG pathway analysis revealed that candidate genes were highly associated with the inflammatory bowel disease, rheumatoid arthritis, toll-like receptor signaling pathway, inflammatory bowel disease, rheumatoid arthritis and toll-like receptor signaling pathway, th17 cell differentiation, osteoclast differentiation and influenza A (Figure 6).

**Figure 5.**
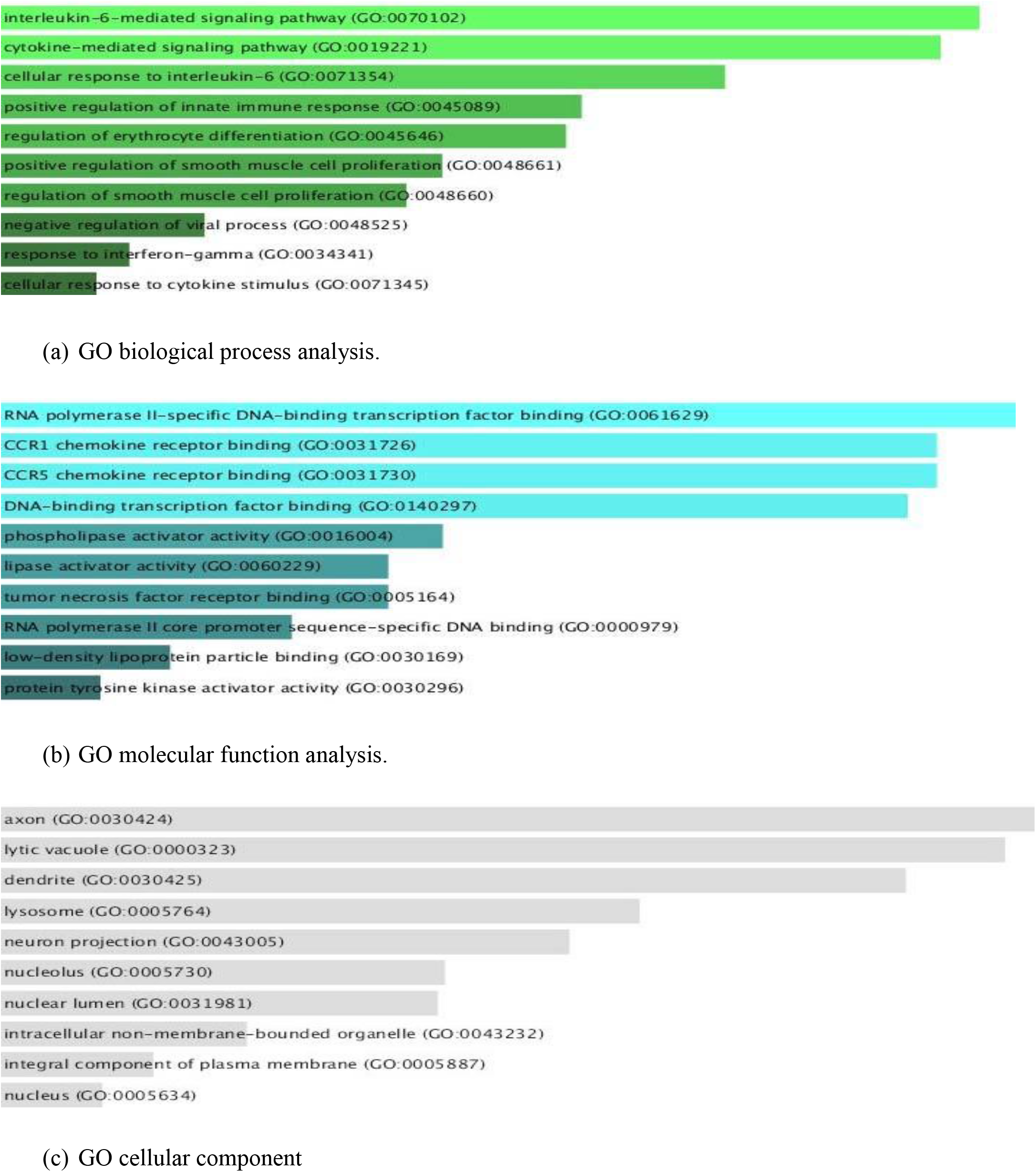
Enriched gene ontology (GO) functions of candidate genes.

**Figure 6.**
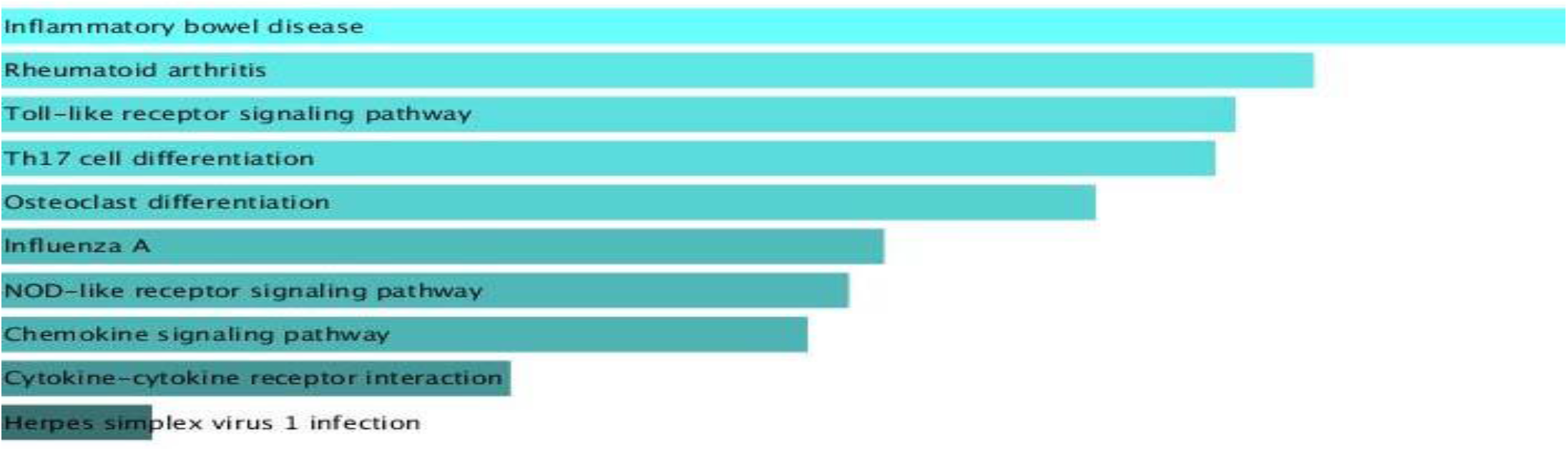
Kyoto Encyclopedia of Genes and Genomes (KEGG) pathway analysis of candidate genes.

**Figure 7.**
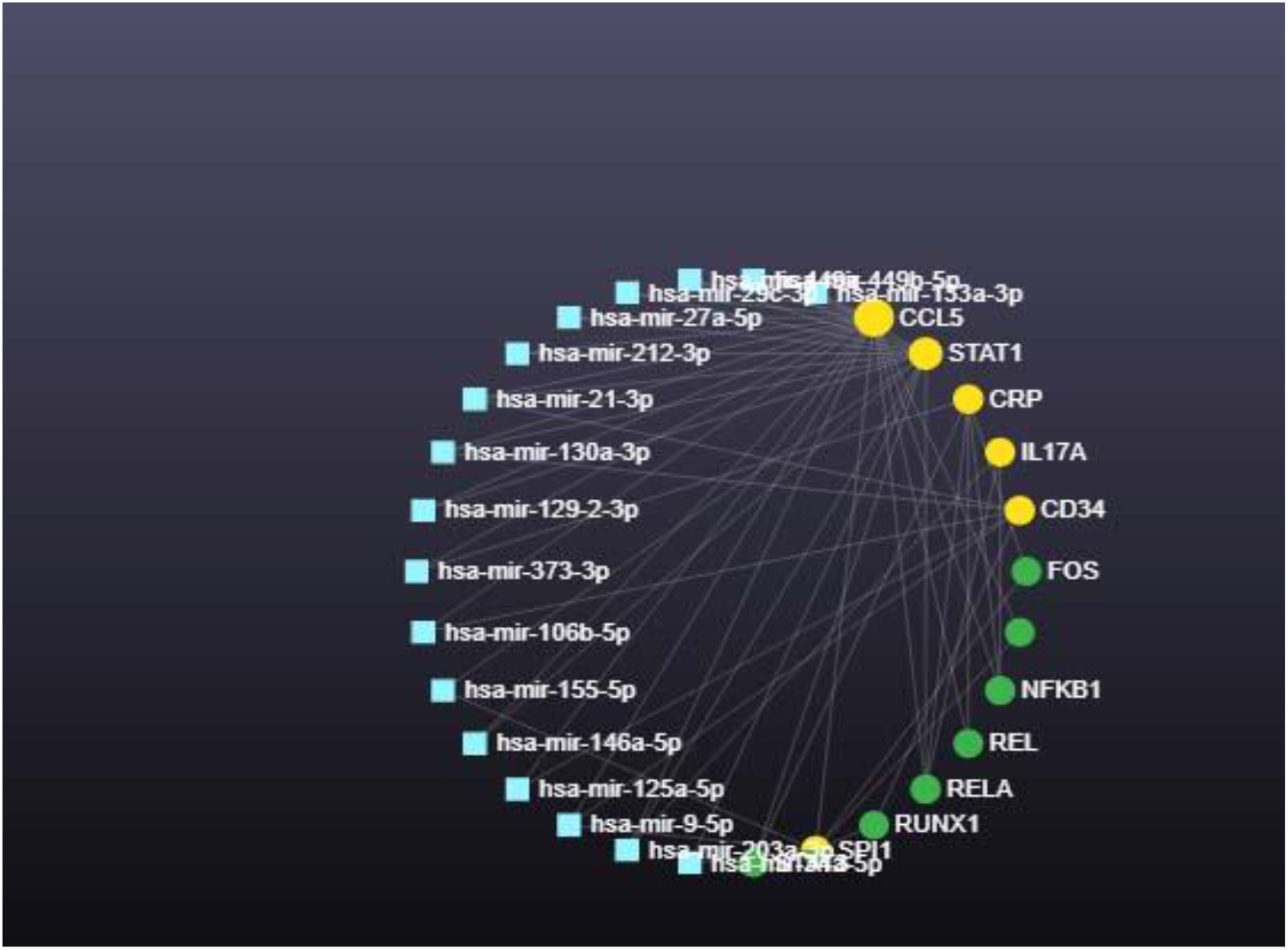
Candidate gene and miRNA interaction graph. The genes, miRNAs, and transcription factor TF are represented in yellow, blue, and green colors respectively.

### 3.4. microRNA interaction analysis

Uploading the candidate genes into the online software generated a network between the genes and miRNAs. 8 transcription factors (TFs) and 17 miRNAs were found associated with the candidate genes. the gene CCL5 has the highest connectivity, with degree score of 20, followed by STAT1 with degree score of 15. SPI1, CRP, CD34, IL17A possess the degree score of 7,6,6,3 respectively.

## 4. Discussion

we identified a total of 1107 significant DEGs between IAV and normal samples, which includes 703 upregulated genes and 404 downregulated genes, conducted a series of analysis to screen candidate genes and pathways related to IAV using bioinformatics tools in this study. Constructing the PPI, 6 candidate genes were identified, CRP, CCL5, IL17A, STAT1, CD34, SPI1 of which 4 of them has been featured in several reports regarding IAV infection and they include CRP, CCL5, IL17A, STAT1.

GO and KEGG analysis investigation conducted to determine functional annotation on candidate gene suggest the role IL6 plays as candidate genes are massively enriched in pathways generating IL6 and other cytokines, while KEGG pathway analysis reveals that candidate genes are enriched in IAV amongst other pathways listed.

The protein encoded by CRP gene belongs to the pentraxin family which also includes serum amyloid P component protein and pentraxin3. Pentraxins are involved in complement activation and amplification via communication with complement initiation pattern recognition molecules, but also complement regulation via recruitment of complement regulators [11]. The short pentraxin SAP has been shown to act as an inhibitor against influenza viruses, binding in a Ca2-dependent manner to mannose-rich glycans on the viral HA to inhibit both hemagglutination and viral infectivity [12] This claim is also corroborated by [13],stating that human and murine pentraxin3 bound to influenza virus and mediated a range of antiviral activities, including inhibition of hemagglutination, neutralization of virus infectivity and inhibition of viral neuraminidase. Antiviral activity was associated with binding of the viral hemagglutinin glycoprotein to sialylated ligands present on pentraxin3. Further in the same report in mouse model experiment, pentraxin3 was seen to be rapidly induced following influenza infection. Like-wise is the therapeutic treatment of mice with human pentraxin3 which resulted in promoting survival and reduced viral load in the lungs following infection with Pentraxin3-sensitive, but not Pentraxin3-resistant, influenza viruses. Indeed, CRP gene can be seen to be involved in the extracellular phase of IAV infection and optimizing this first phase of immune response may be crucial in mitigating further viral hold that may be fatal especially before it gets into the cell for its imminent replication.

The CCL5 gene is one of many chemokine genes. Chemokines are involved in immunoregulatory and inflammatory processes.as part of the CC subfamily, they function as a chemoattractant for blood monocytes, memory T helper cells and eosinophils. This chemokine functions as a natural ligand for the chemokine receptor CCR5 [14]. Ligands of CCR5 receptor play a key role among the molecules enhanced in the infected respiratory tract as they triggers the migration of inflammatory cells to the alveoli [15]. This process is triggered by IAV-induced-cytokine storm; the protective antiviral effects promoted by the interferon (IFN) are overwhelmed by other cytokines, like TNF-α and IL-6. At the infection site at the same time, the expression of adhesion molecules is increased and chemotactic factors which includes CCL3, CCL4 and CCL5, are released, attracting leukocytes and monocytes into the alveolar lumen [16]. The binding of chemokines CCL3, CCL4 and CCL5 to the CCR5 receptor allows participation in leukocyte activation, chemotaxis, cytokine secretion and cell proliferation [17], which consequently may result into tissue damage and respiratory distress.

The IL17A gene belongs to the IL-17 receptor family (IL-17RA-E), consisting of five members, involved in coding for pro-inflammatory cytokine proteins by activated T cells. Production of inflammatory chemicals, chemokines, antimicrobial peptides, and remodeling proteins are all induced by IL-17A-mediated downstream pathways [18]. IL-17A) has been proven to play crucial roles not only in autoimmune disorders but also in viral infectious diseases as its appropriate(regulated) levels can trigger the expression of a series of cytokines, chemokines, antimicrobial peptides, IL-6, G-CSF, TNF-α and CXCL1, to kill pathogens [19].However, overexpression of IL17A may result in severe inflammation and disease progression due to excessive recruitment and accumulation of inflammatory cells causing serious lung injury as shown in a recent study in mice infected with IAV for 2 day [20], perceived as autoimmune disorder. Considering the mode of action in the functionality of this gene, regulation or complete knockout of IL17A gene might be a potential approach for circumventing the abnormal immune response during IAV infection.

The STAT1 gene is a member of the STAT protein family. they act as transcription activators after being phosphorylated by receptor associated kinases in response to cytokines and growth factors. The protein encoded by this gene can be activated by various ligands including interferon-alpha, interferon-gamma, EGF, PDGF and IL6 [21] in response to viral infection detected by the pattern recognition receptors. This protein mediates the expression of a variety of genes, which is thought to be important for cell viability in response to pathogenic invasion and diverse cell stimuli. The protein plays an important role in immune responses to viral, fungal and mycobacterial pathogens. STAT1 is reported to be known as an important anti-viral molecule during IAV infection [22], this claim also corroborated by reports conducted from in vitro and in vivo studies on STAT1 knock-out mice; STAT1-deficiency could accelerate the pathogenesis of IAV which lack the STAT1 protein thereby indicating the role of STAT1 in regulating IAV replication and inflammatory response [23].

## 5. Conclusions

Discovering the possibility of enhancing the functionality of innate immune response by the use of antiviral agents in preparedness on the event of another outbreak of IAV cannot be overemphasized. This process begins by identification, followed by comprehensive understanding of the pathways and mechanisms of action of key proteins involved in innate immune response. This study tried to identify some human innate immune response genes and pathway regulatory related to IAV infection by a series of bioinformatics analysis of DEGs between IAV samples and normal samples, as findings of this study may help us understand the underlying molecular mechanisms of the innate immune response genes. Genes such as, CRP, CCL5, IL17A and STAT1 showed potentials worthy of further analysis in IAV studies.

## Acknowledgments (All sources of funding of the study must be disclosed)

This research has no acknowledgement

